# Antenatal Screening for Group B Streptococcus in the Setting of Preterm Premature Rupture of Membranes: Empiric versus Culture-based Prophylaxis

**DOI:** 10.1101/201400

**Authors:** Leena B. Mithal, Nirali Shah, Anna Romanova, Emily S. Miller

## Introduction

Group B streptococcus (GBS) bacteria colonizes the genital tract in 10-30% of pregnant women.^1,2^ GBS colonization during pregnancy is the primary risk factor for neonatal GBS infection with its accompanying risks of significant infant morbidity and mortality.^3-5^ Over the last several decades, intrapartum intravenous administration of antibiotics to women at risk for transmitting GBS has greatly reduced neonatal disease.^3,6,7^ Current CDC guidelines recommend routine screening for GBS at 35-37 weeks gestation and subsequent administration of intrapartum antibiotics if the GBS screen is positive (universal culture-based prophylaxis). However, this approach is complicated in the setting of preterm premature rupture of membranes (PPROM), which is causes up to 30% of preterm births.^8^ Prolonged rupture of membranes (>12 hours) is associated with increased risk of transmission of GBS to the newborn.^9^ Furthermore, preterm infants are at increased risk for GBS infection, early-onset neonatal sepsis (EONS), and severe infection with complications, particularly at <34 weeks gestational age (GA).^5,10,11^

Screen-based intrapartum antibiotic prophylaxis (IAP) has been shown to be superior compared to treatment based on risk factors, but there is no adequate data regarding reliability of GBS screening in the setting of PPROM.^12^ Women with PPROM at <33 weeks gestation will not yet typically have undergone routine outpatient GBS screening. In this case, the recommendation is to screen for GBS on admission, prior to receipt of any antibiotics. Upon the diagnosis of labor or at the time of induction, the GBS status, if available, should guide whether or not to use intravenous intrapartum antibiotics for GBS (i.e., a culture-based IAP approach).^13^ If culture results are not available, empiric IAP is provided.^3,14,15^

There are data in term and preterm deliveries that indicate that errors at multiple points of care lead to suboptimal recommended treatment of GBS colonization in pregnant women prior to delivery, specifically limited GBS screening on hospital admission and limited administration of IAP.^16^ Furthermore, in cases of neonatal GBS infections, up to 60-80% of women had a negative GBS screen.^17^ Thus, the limitations in sensitivity of GBS culture also results in insufficient IAP for some women.^7^ Given the concerns for possible under-detection of GBS colonization and the increased morbidity and mortality associated with GBS infection in preterm neonates, it is currently unclear what the infectious morbidity is to neonates born to GBS-screen negative mothers. Therefore, the objective of this study was to evaluate if empiric IAP (administered universally to women with PPROM) is associated with reduced risk of EONS and neonatal GBS infection compared to culture-based IAP in the setting of PPROM.

## Methods

### Study design and subjects

We conducted a retrospective cohort study of women who delivered at Northwestern Memorial’s Prentice Women’s Hospital, a tertiary care hospital in Chicago, IL between January 2010 and October 2015. Women were included if they were at least 18 years of age, carrying a singleton gestation, and were diagnosed with PPROM between 23-33 weeks. Mothers with suspected fetal anomalies were excluded.

Routine clinical care of women presenting with PPROM during the study period, in accordance with the American College of Obstetricians and Gynecologists (ACOG) guidelines^15^ included clinical confirmation of the diagnosis and assessment of fetal well-being. Antenatal betamethasone was administered and, if labor had not ensued, latency antibiotics were initiated. These consisted of ampicillin and erythromycin for a total of seven days. Women were started on magnesium for fetal neuroprotection and were monitored on the Labor and Delivery unit for twelve hours. Thereafter, if maternal and fetal stability remained, she was transferred to the inpatient antepartum unit for close monitoring. Delivery was recommended if there was concern for chorioamnionitis or a clinically significant abruption. Otherwise an induction was planned for 34 weeks gestation. Rescue steroids were reserved for documentation of a clinical change suggestive of delivery within seven days.

Comprehensive clinical data were abstracted from maternal and infant electronic medical records by authorized research personnel. Maternal data included sociodemographics, past medical history, past obstetrical history, labor and delivery outcomes including length of antepartum admission, indication for delivery, development of fever or chorioamnionitis, and laboratory findings including GBS screening culture. Data on maternal antibiotic administration was abstracted, including GBS IAP and administration of latency antibiotics, for which standard of care was to administer ampicillin and erythromycin for seven days.^4^

Neonatal data were obtained from linked chart within the electronic medical record system. Neonatal data included one and five minute Apgar scores, temperature abnormalities (recorded as the highest and the lowest temperature), clinical diagnosis of sepsis, any infectious workup results (e.g. CBC with differential, C-reactive protein (CRP), blood and cerebrospinal fluid cultures), antibiotic administration, and neonatal intensive care unit hospital length of stay.

### Infant outcomes

The primary endpoint was EONS. Based on culture results, antibiotic treatment and laboratory data, patients were classified as EONS according to pre-determined criteria based on upon National Institute of Child Health and Human Development definitions for premature infants.^11^ EONS was defined as having received an antibiotic treatment course of ≥5 days starting within 72 hours of life and either 1) a positive bacterial blood culture ≤72 hours of life or 2) other abnormal laboratory results indicative of infection. Abnormal laboratory results indicative of infection were: CRP ≥1mg/dL^18^, absolute neutrophil count (ANC) outside normal range for GA^19,20^, and immature-to-total (I:T) neutrophil ratio >0.22^21^. Analyses were done using one and two abnormal laboratory indices as criteria for presumed EONS. The secondary neonatal outcome measure was early onset neonatal GBS infection, defined as GBS bacteremia and/or meningitis.

### Statistical analyses

Bivariate analysis of demographics, risk factors, and outcomes were conducted using Chi-squared or Fisher exact tests for categorical variables and Student’s t or Wilcoxon rank sum tests for continuous variables. Multivariable analysis was performed to study the association between IAP strategy and EONS/GBS infection. P-values of <0.05 were considered statistically significant. STATA^®^ software, version 13.1 (College Station, TX) was used for this analysis.

### Ethics statement

The study was approved by the Northwestern University Feinberg School of Medicine Institutional Review Board prior to its initiation. Investigations were conducted according to the principles expressed in the Declaration of Helsinki.

## Results

### Maternal data

There were 270 maternal/infant pairs that met inclusion criteria. Culture-based IAP was performed in 136 (50.4%) women and the remaining 134 women were given empiric IAP. Of women with culture-based IAP strategy, 36 (26.5%) were GBS positive and received antibiotics during labor and delivery. The remaining 100 did not receive IAP towards GBS prophylaxis. Table 1 shows maternal characteristics. There were no differences in advanced maternal age, race/ethnicity, parity, history of a prior preterm birth, body mass index (BMI) at delivery, tobacco use, or diabetes, however, diabetes was on threshold of significance with 11.8% in culture-based IAP group versus 5.2% in empiric IAP group (p=0.05).

**Table 1.**
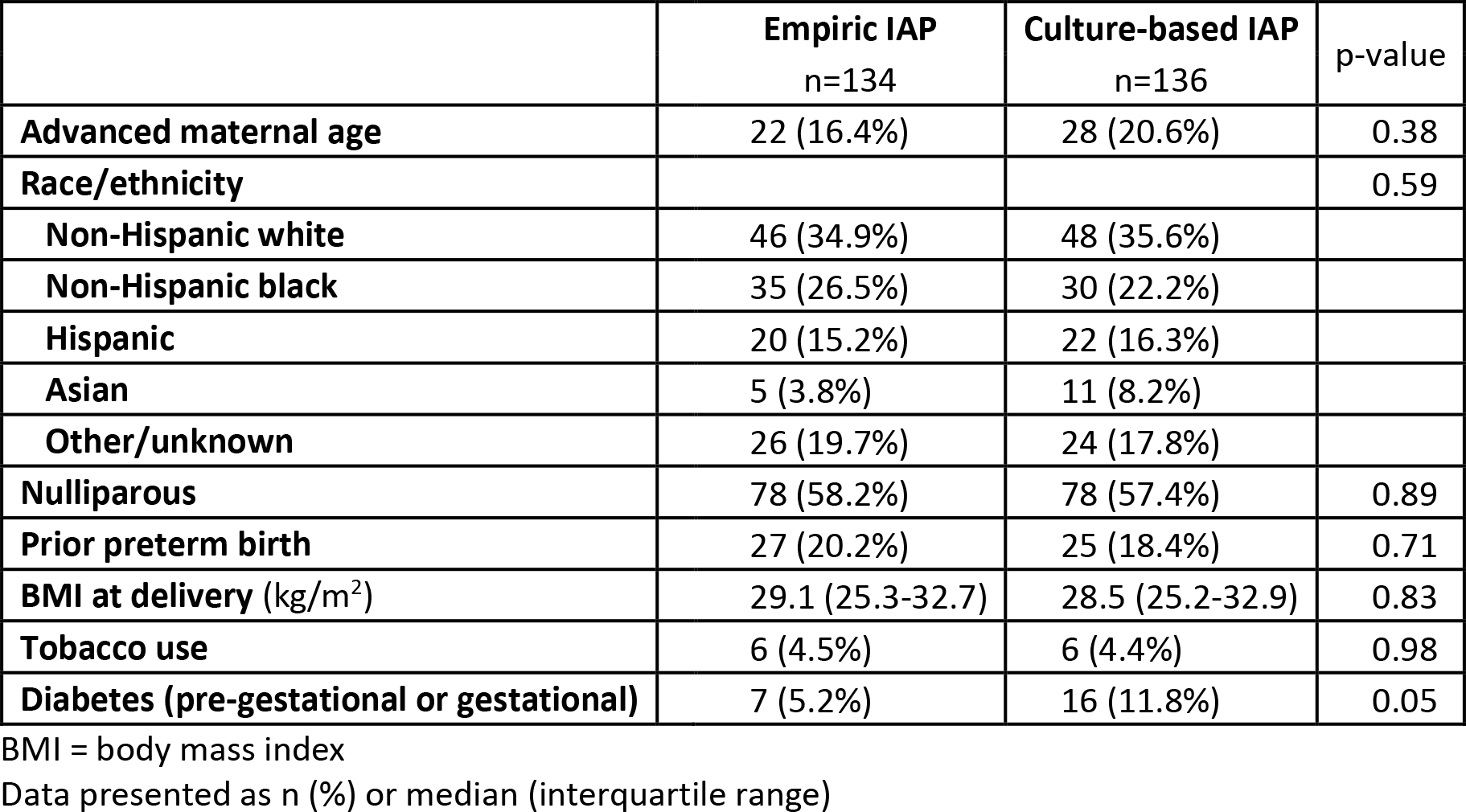
Maternal characteristics stratified by intrapartum antibiotic prophylaxis strategy

|  | **Empiric IAP**<br>n=134 | **Culture-based IAP**<br>n=136 | **p-value** |
| --- | --- | --- | --- |
| **Advanced maternal age** | 22 (16.4%) | 28 (20.6%) | 0.38 |
| **Race/ethnicity** |  |  | 0.59 |
| **Non-Hispanic white** | 46 (34.9%) | 48 (35.6%) |  |
| **Non-Hispanic black** | 35 (26.5%) | 30 (22.2%) |  |
| **Hispanic** | 20 (15.2%) | 22 (16.3%) |  |
| **Asian** | 5 (3.8%) | 11 (8.2%) |  |
| **Other/unknown** | 26 (19.7%) | 24 (17.8%) |  |
| **Nulliparous** | 78 (58.2%) | 78 (57.4%) | 0.89 |
| **Prior preterm birth** | 27 (20.2%) | 25 (18.4%) | 0.71 |
| **BMI at delivery (kg/m<sup>2</sup>)** | 29.1 (25.3-32.7) | 28.5 (25.2-32.9) | 0.83 |
| **Tobacco use** | 6 (4.5%) | 6 (4.4%) | 0.98 |
| **Diabetes (pre-gestational or gestational)** | 7 (5.2%) | 16 (11.8%) | 0.05 |

BMI = body mass index
Data presented as n (%) or median (interquartile range)

Details of the antenatal management and delivery were also largely similar between groups including receipt of a complete course of antenatal steroids, preterm labor, route of delivery, and need for emergent delivery. As anticipated, the duration of latency was longer in the culture-based IAP compared to the empiric IAP and, accordingly, the gestational age at delivery was longer in the culture-based IAP compared to the empiric IAP groups. Sixty-four percent of women in the cohort were on latency antibiotics at time of delivery and 87% of women in the cohort received antibiotics within 7 days of delivery; 95% in empiric IAP versus 80% in culture-based IAP group (p<0.01). Mothers in the culture-based IAP group were significantly more likely to undergo an induction of labor (p=0.02). Infant sex and birthweight were similar between groups.

**Table 2.** Antenatal and delivery data stratified by intrapartum antibiotic prophylaxis strategy

|  | **Empiric IAP**<br>n=134 | **Culture-based IAP**<br>n=136 | **p-value** |
| --- | --- | --- | --- |
| **Antenatal steroids** | 108 (80.6%) | 120 (88.2%) | 0.08 |
| **Induction of labor** | 17 (12.7%) | 32 (23.5%) | **0.02** |
| **Preterm labor** | 92 (68.7%) | 81 (59.6%) | 0.12 |
| **Cesarean delivery** | 40 (29.9%) | 40 (29.4%) | 0.94 |
| **Emergent delivery** | 25 (18.7%) | 16 (11.8%) | 0.12 |
| **Chorioamnionitis** | 14 (10.5%) | 25 (18.4%) | 0.06 |
| **Gestational age, weeks** | 31.6 (29.4-33.0) | 32.3 (29.6-33.7) | **0.01** |
| **Infant sex (M)** | 69 (51.5%) | 63 (46.3%) | 0.40 |
| **Birthweight** | 1695 ± 516 | 1772 ± 540 | 0.24 |

Data presented as n (%), median (interquartile range), or mean (standard deviation)

### Infant outcomes

Nine (3.3%) infants in the cohort had bacteremia in the first 72 hours of life treated with antibiotics. The organisms isolated in blood culture from infants of mothers in the empiric IAP group were *E coli* (n=2), GBS (n=1), *Haemophilus influenzae* (n=1) and *Klebsiella pneumoniae* (n=1). Bacteremia in the culture-based IAP group infants were *E coli* (n=2), GBS (n=1), and coagulase negative staph species (n=1). There was no significant difference in the risk of either early-onset GBS infection or bacteremia between groups (Table 3). Abnormal laboratory results indicative of infection were also not significantly different between groups, in isolation. We analyzed two grades of strictness for presumed EONS by antibiotics and laboratory criteria. There was no increased risk of EONS by bacteremia and/or either one or two accessory laboratory criteria. These findings persisted after controlling for potential confounders (Table 3). Early life antibiotic treatment of infants did not differ between group, specifically infants receiving no antibiotics, <5 days of antibiotics (a “rule-out” of EONS), and ≥5 days of antibiotics (an antibiotic treatment course for proven or presumed EONS), as shown in Figure 1. A post-hoc power calculation demonstrated that this study had 90% power to detect at least a 2-fold difference in EONS between the two IAP strategies.

**Table 3.** Neonatal outcomes following empiric versus culture-based maternal IAP

|  | Empiric IAP<br>n=134 | Culture-based IAP<br>n=136 | p-value | aOR* | 95% CI |
| --- | --- | --- | --- | --- | --- |
| **Neonatal bacteremia** | 5 (3.7%) | 4 (2.9%) | 0.748 | 0.91 | 0.23-3.57 |
| **GBS bacteremia** | 1 (0.7%) | 1 (0.8%) | 1 | N/A | N/A |
| **Antibiotics ≥5 days** | 44 (32.8%) | 42 (30.9%) | 0.73 | 1.11 | 0.63-1.98 |
| **Absolute Neutropenia** | 40 (29.9%) | 40 (29.4%) | 0.937 | 0.97 | 0.57-1.65 |
| **Elevated I:T ratio** | 31 (23.1%) | 25 (18.4%) | 0.336 | 0.82 | 0.45-1.51 |
| **Abnormal C-reactive protein** | 31 (23.1%) | 32 (23.5%) | 0.936 | 1.29 | 0.65-2.58 |
| **EONS (culture or 2 abnl labs)** | 24 (17.9%) | 15 (11%) | 0.108 | 0.66 | 0.31-1.39 |
| **EONS (culture and/or 1 abnl lab)** | 37 (27.6%) | 30 (22.1%) | 0.291 | 0.86 | 0.47-1.58 |

\* Adjusted for indication for delivery and GA

**Figure 1.**
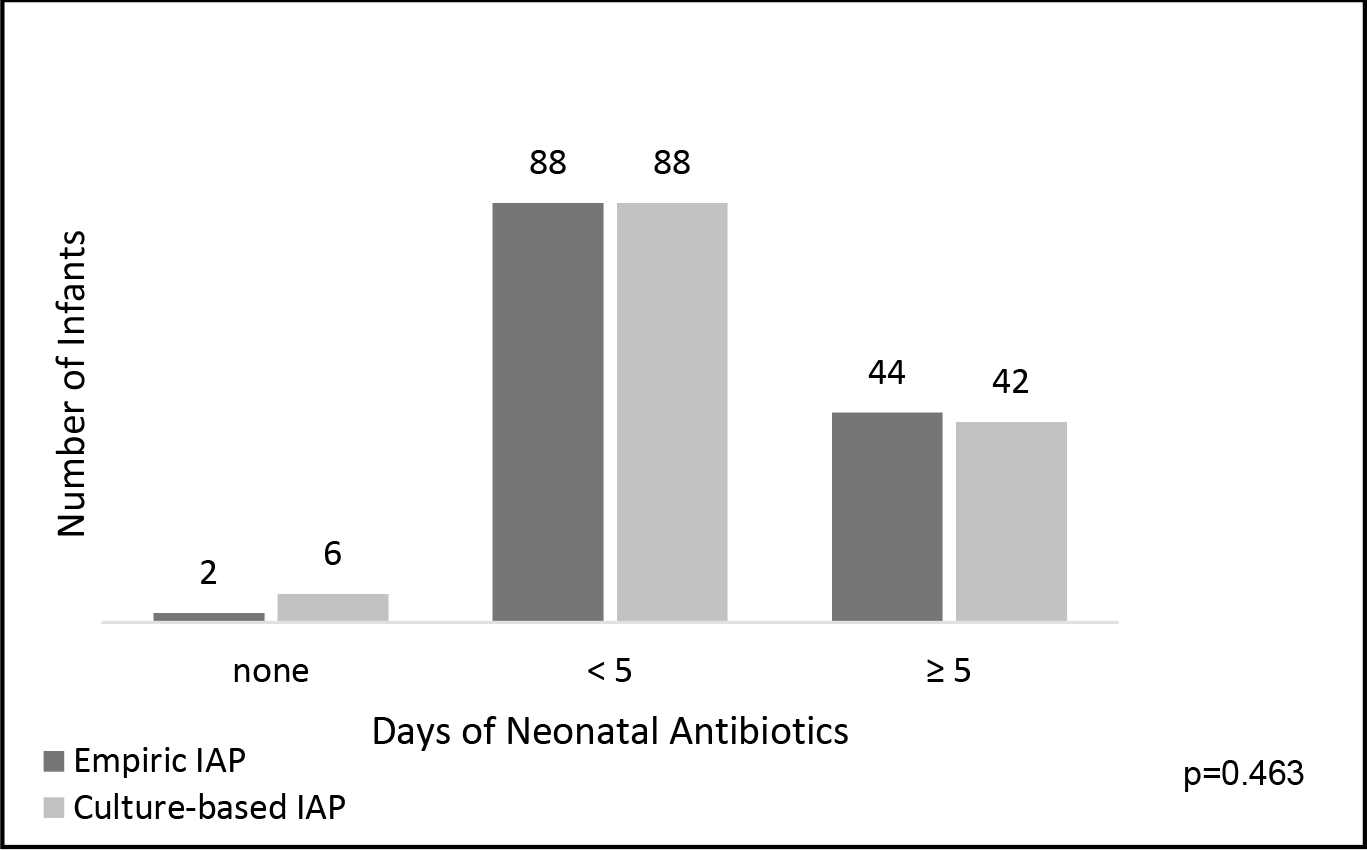
Days of early life neonatal antibiotic administration. Figure 1 legend. Number of infant antibiotic days, with course starting in first 72 hours of life (early life antibiotics).

## Discussion

This study reports and compares EONS and neonatal early-onset GBS infection in infants of mothers with PPROM prior to 33 weeks who received culture-based IAP versus those who received empiric IAP (without GBS screening culture sent or resulted). In our retrospective analysis, we found no difference in EONS or GBS infection between these IAP strategies. This study provides evidence supporting the current recommended guidelines for culture-targeted prophylaxis, even for women with PPROM and resultant preterm delivery.

The appropriate use of antibiotics in women with PPROM and high likelihood of preterm delivery is important for both maternal and infant outcomes. There are risks and benefits to widespread, empiric antibiotic use, certainly administered directly to the infant, but also to mother prior to delivery. Risks include an increased risk of necrotizing enterocolitis in the infant,^22,23^ and recent ongoing research suggests that maternal antibiotics prior to delivery are associated with notable alternations in an infant’s early life microbiome diversity with potential far-reaching effects on an infant’s metabolic, immune, and allergic health.^24^ However, administration of certain antibiotics have been shown to prolong latency,^15,25,26^ and there is clearly a significant decrease in GBS disease since institution of GBS IAP recommendations.

To date, there has been a paucity of data on the optimal strategy for IAP administration for women with PPROM, who do not have GBS culture results available at the time of rupture. The theoretical possibility of increased risk of neonatal infection given imperfect sensitivity of culture and adherence to screen, result, and antibiotic recommendations led to this retrospective study. The negative predictive value of GBS culture declines outside a 5 week window between culture and delivery, thus early screening would not address this problem. Our study reinforces that screening culture and culture-result based IAP decisions are not causing a missed opportunity for EONS prevention. Although our study was retrospective and EONS is relatively infrequent, our results show actually a higher percent of EONS cases present in the empiric IAP patients than culture-based IAP. This may represent a higher rate of spontaneous preterm labor versus induced delivery in the culture-based IAP group, resulting in time for culture results to be obtained. GBS infection was not different between groups, which is the primary objective of the CDC’s IAP guidelines.

The majority of women in both groups received antibiotics to promote gestational latency which remains standard clinical practice. The effectiveness of latency antibiotics >7 days from delivery versus active treatment with latency antibiotics in the peripartum period on EONS and GBS infection is unknown. Latency antibiotics could play a confounding or interaction role in the association between IAP and EONS, and thus requires further study.

The strengths of this study include detailed maternal and infant data on demographics, risk factors, antibiotics, and infant outcomes for a cohort of maternal-infant dyads. The granular definitions of EONS, incorporating both proven and presumed sepsis, microbiology results, and clinical antibiotic treatment decision strengthens the analysis of neonatal outcomes. A notable limitation in this study is the few infants with proven early-onset GBS infection, and culture-proven infection in general which limits our power to detect potentially clinically important differences in this relatively rare, but highly morbid, condition.

In conclusion, this study demonstrates that a screen and culture-based IAP strategy for women presenting with PPROM at <33 weeks appears to be appropriate, without obvious missed opportunities for neonatal infection prevention and equivalent overall outcomes. Further data on timing of culture-based screening, other antibiotic treatments for PPROM management, and larger prospective observational studies are necessary to advance perinatal care and optimize targeted antibiotic exposures to mother and infant.

## Acknowledgements

We would like to acknowledge Northwestern Medicine Enterprise Data Warehouse, funding support from NICHD K12 HD050121-09 (ESM), and the patients at Prentice Women’s Hospital who make this study possible.

